# Mfge8 attenuates human gastric antrum smooth muscle contractions

**DOI:** 10.1101/2020.09.09.289173

**Authors:** Wen Li, Ashley Olseen, Yeming Xie, Cristina Alexandru, Brian A. Perrino

## Abstract

Coordinated gastric smooth muscle contraction is critical for proper digestion and is adversely affected by a number of gastric motility disorders. In this study we report that the secreted protein Mfge8 (milk fat globule-EGF factor 8) inhibits the contractile responses of human gastric antrum muscles to cholinergic stimuli by reducing the inhibitory phosphorylation of the MYPT1 (myosin phosphatase-targeting subunit 1) subunit of MLCP (myosin light chain phosphatase), resulting in reduced LC20 (smooth muscle myosin regulatory light chain 2) phosphorylation. We show that endogenous Mfge8 is bound to its receptor, α8β1 integrin, in human gastric antrum muscles, suggesting that human gastric antrum muscle mechanical responses are regulated by Mfge8. The regulation of gastric antrum smooth muscles by Mfge8 and α8 integrin functions as a brake on gastric antrum mechanical activities. Further studies of the role of Mfge8 and α8 integrin in regulating gastric antrum function will likely reveal additional novel aspects of gastric smooth muscle motility mechanisms.

## Introduction

Digestion of ingested food by the stomach involves accommodation, chemical and mechanical disruption of solids into chyme, and controlled emptying into the duodenum. To carry out these functions, the stomach is comprised of functional anatomic regions with distinct motility patterns [1, 2]. The fundus relaxes to accommodate ingested food and then tonically contracts to move the contents into the distal stomach where the solids are reduced in size by peristaltic contractions. Gastric emptying is regulated by contractions of the antrum and the resistance provided by the pyloric canal. Healthy gastric function depends on properly coordinated motor activities of the proximal and distal stomach [3]. Animal models have been studied for many years, but the regulatory mechanisms underlying the motor activities of the human stomach are not as well understood [4, 5].

Membrane depolarization of gastrointestinal (GI) smooth muscles triggers contraction by opening voltage-dependent (L-type) Ca^2+^ channels, non-selective cation currents, and other mechanisms that contribute to the Ca^2+^ influx and the increase in [Ca^2+^]_i_ [6, 7]. The increase in [Ca^2+^]_i_ activates calmodulin-dependent myosin light chain kinase (MLCK) to phosphorylate LC20 at S19 (pS19), stimulating myosin ATPase activity to generate cross-bridge cycling and contraction [8, 9]. Termination of the contractile signal decreases [Ca^2+^]_i_ by Ca^2+^ removal mechanisms, and inactivation of MLCK [10, 11]. LC20 is then dephosphorylated by MLCP, leading to relaxation [12, 13]. MLCP activity is inhibited by upstream kinase-dependent signaling pathways [14-16]. Phosphorylation of the protein kinase C-(PKC) potentiated phosphatase inhibitor protein-17 kDa (CPI-17) by PKC greatly increases its inhibition of MLCP [17, 18]. Phosphorylation of MYPT1 at T696 (human isoform numbering) inhibits MLCP activity [19, 20]. Phosphorylation of MYPT1 T853 by Rho-associated coiled-coil protein kinase 2 (ROCK2) reduces the affinity of MLCP to myosin filaments in vitro [21]. However, ROCK2 phosphorylation of MYPT1 T853 does not appear to affect MLCP activity in vivo [22, 23]. In addition, expression of the MYPT1 T853A mutant does not affect agonist-induced LC20 phosphorylation and force development in bladder and ileum smooth muscles [22, 24]. Thus, although it is elevated by ROCK2 activation, MYPT1 T853 phosphorylation is not necessary for agonist-induced Ca^2+^ sensitization of smooth muscle [22-24]. However, ROCK2 activity in smooth muscles is clearly required for Ca^2+^ sensitization and augmented contraction [22]. Therefore, the level of MYPT1 T853 phosphorylation can be used as an indicator of myofilament Ca^2+^ sensitization in smooth muscles. Inhibiting MLCP while activating MLCK generates greater force by further increasing LC20 phosphorylation [25, 26]. This phenomenon was termed “Ca^2+^ sensitization of the contractile apparatus,” to describe the increased Ca^2+^sensitivity of the contractile response [9].

A novel mechanism regulating ROCK2-dependent myofilament Ca^2+^ sensitization in gastric smooth muscles has recently been described in murine gastric antrum muscles, involving the secreted protein Mfge8 [27]. The binding of Mfge8 to α8β1 integrin heterodimers results in the inhibition of MYPT1 phosphorylation by ROCK2 and inhibition of antral contractility and gastric emptying [27]. In contrast, in Mfge8^-/-^ mice, or α8 integrin^-/-^ mice, MYPT1 phosphorylation and antral contractility and gastric emptying are increased [27]. These findings indicate that Mfge8 binding to α8β1 integrins acts as a “brake” on gastric muscle contractions, and more importantly, suggest that disrupting the binding of Mfge8 to α8β1 integrins in gastric smooth muscles improve or restore gastric motility in patients with gastroparesis. We have previously found that MYPT1 T853 is constitutively phosphorylated in human gastric smooth muscles, and is decreased by ROCK2 inhibition [28-30]. However, whether Mfge8 regulates MYPT1 phosphorylation and the contractile responses of human gastric smooth muscles has not been reported. In this report, we show that, similar to mouse gastric antrum muscles, Mfge8 is present in human gastric antrum muscles and is constitutively bound to α8β1 integrin. We also show that exogenously added Mfge8 inhibits the contractions evoked by electric field stimulation of cholinergic motor neurons, and the contractile responses to the cholinergic agonist carbachol (CCh), and decreases the phosphorylation of MYPT1 T696 and T853 and LC20 S19 in human gastric antrum muscles.

## Materials and Methods

### Human stomach smooth muscles

The use of human resected stomach tissues was approved by the Human Subjects Research Committees at the Renown Regional Medical Center and the Biomedical Institutional Review Board at the University of Nevada, Reno, and was conducted in accordance with the Declaration of Helsinki (revised version, October 2008, Seoul, South Korea). All patients provided written informed consent. Resected stomach specimens were acquired immediately after surgery from patients undergoing vertical sleeve gastrectomy. The resected stomach tissue was placed into ice-cold Krebs–Ringer buffer (KRB; composition (in mM): NaCl 118.5, KCl 4.5, MgCl_2_ 1.2, NaHCO_3_ 23.8, KH_2_PO4 1.2, dextrose 11.0, and CaCl_2_ 2.4; for transport to the laboratory. The gastric fundus region was identified by its bulbous appearance, and the gastric antrum region was identified by its narrow tapered shape. The resected stomach tissues were opened along the staples, laid out flat, and pinned to a Sylgard-lined dish containing oxygenated KRB. The mucosa and submucosa were removed by sharp dissection. Gastric antrum muscles were mapped and obtained from regions 13–16 [31]. Rectangular strips (∼4 mm × 10 mm × 2 mm) of full thickness muscle were used for the contractile studies and the protein phosphorylation studies.

### Mechanical responses

Gastric antrum smooth muscle strips were attached to a Fort 10 isometric strain gauge (WPI, Sarasota, FL, USA), in parallel with the circular muscles, and pretreated with 2 µM neostigmine for 10 min at 37°C in oxygenated KRB, and three 1 min washes with KRB, to remove any residual curariform neuromuscular paralytics [32]. Contractions were measured in static myobaths with oxygenated Krebs bubbled with 97% O_2–_3% CO_2_ at 37°C, the pH of KRB was 7.3–7.4). Each strip was stretched to an initial resting force of ∼0.8 g and then equilibrated for 45 min-60 min in 37°C oxygenated KRB. To measure the contractile responses to KCl or CCh, the muscle strips were incubated with 0.3 µM tetrodotoxin to eliminate motor neuron activity. To measure contractile responses in response to electrical field stimulation, the muscle strips were incubated with LNNA and MRS2500 to eliminate nitrergic and purinergic motor neuron activity [30]. Contractile activity was acquired and analyzed with AcqKnowledge 3.2.7 software (BIOPAC Systems, www.biopac.com).

### Automated capillary electrophoresis and immunodetection with Wes Simple Western

For automated capillary electrophoresis and Western blotting by Wes, the muscles were submerged into ice-cold acetone/10 µM dithiothreitol (DTT)/10% (w/v) trichloroacetic acid for 2 min, snap-frozen in liquid N_2_, and stored at −80°C for subsequent Wes analysis [32, 33]. Muscles were washed in ice-cold-acetone–10 µM DTT for 1 min, 3 times, followed by a 1 min wash in ice-cold lysis buffer (mM: 50 Tris–HCl pH 8.0, 60 β-glycerophosphate, 100 NaF, 2 EGTA, 25 sodium pyrophosphate, 1 DTT, 0.5% NP-40, 0.2% sodium dodecyl sulfate and protease inhibitors [28]. Tissues were homogenized in 0.5 ml lysis buffer in a Bullet Blender (0.01% anti-foam C, one stainless steel bead per tube, speed 6, 5 min), then centrifuged at 16,000 x g, for 10 min at 4°C. Supernatants were stored at −80°C. Protein concentrations of the supernatants were determined by the Bradford assay using bovine γ-globulin as the standard. Protein expression and phosphorylation levels were measured and analyzed according to the Wes User Guide using a Wes Simple Western instrument from ProteinSimple (www.proteinsimple.com). The protein samples were mixed with the fluorescent 5X master mix (ProteinSimple) and then heated at 95°C for 5 min. Boiled samples, biotinylated protein ladder, blocking buffer, primary antibodies, ProteinSimple horseradish peroxidase-conjugated anti-rabbit or anti-mouse secondary antibodies, luminol-peroxide and wash buffer were loaded into the Wes plate (Wes 12–230 kDa Pre-filled Plates with Split Buffer, ProteinSimple). The plates and capillary cartridges were loaded into the Wes instrument, and protein separation, antibody incubation and imaging were performed using default parameters. Compass software (ProteinSimple) was used to acquire the data, and to generate image reconstruction and chemiluminescence signal intensities. The protein and phosphorylation levels are expressed as the area of the peak chemiluminescence intensity. The following primary antibodies were used for Wes analysis: mouse anti-integrin-α8, MAB6194, www.rndsystems.com; rabbit anti-integrin-β1, sc-8978; rabbit anti-LC20, sc-15370; www.scbt.com; rabbit anti-Mfge8, HPA002807, www.sigmaaldrich.com; rabbit anti-MYPT1 (PPP1R12A), sc-25618; rabbit anti-pT696-MYPT1, sc-17556-R; rabbit anti-pT853-MYPT1, sc-17432-R; rabbit anti-pS19-LC20, PA5-17726, www.thermofisher.com.

### Immunofluorescence and in situ proximity ligation assay (isPLA)

For both immunofluorescence and isPLA the gastric antrum smooth muscle strips were fixed with 4% paraformaldeyde in PBS, and then cryo-protected with PBS/30% sucrose at 4°C, embedded in OCT, and frozen at −80°C [34]. The blocks were cut using a microtome into 10 µm sections and placed onto Vectabond (SP-1800) coated glass slides (Fisherbrand Superfrost Plus Microscope Slides, 12-550-15). After 20 min microwave heat-induced antigen retrieval in Tris-EDTA buffer (10 mM Tris base, 1 mM EDTA solution, 0.05% Tween 20, pH 9.0), the slides were permeabilized and blocked with PBS containing 0.2% Tween-20 and 1% BSA for 10 min at room temperature. The slides were then incubated overnight at 4°C with the appropriate primary antibody as indicated below. Immunofluorescent labeling was performed with the appropriate Alexa-488 or Alexa-594 conjugated secondary antibody (Cell Signaling Technology, www.cellsignal.com) against the primary antibody (1:500 for 30 min at room temperature in PBS). isPLA was performed according to the manufacturer’s instructions using the Duolink In Situ Detection Reagents Red DUO92008 (Sigma-Aldrich, Olink Bioscience, Sweden, www.sigmaaldrich.com) [34]. The muscle sections were incubated with each primary antibody (1:400 dilution) sequentially for 1 h at room temperature. The slides were then incubated with the appropriate PLA probes (diluted 1:5 in PBS containing 0.05% Tween-20 and 3% bovine serum albumin) in a pre-heated humidified chamber at 37°C for 1 h, followed by the ligation (30 min, 37°C) and amplification (100 min, 37°C) reactions. Mounting medium with DAPI was used to label nuclei blue. It has been reported that the number of PLA signals can decrease as kits get older [35]. We did not experience any differences in the PLA results as the kits aged. However, control and treated muscle sections were compared using Duolink Detection kits from the same lot number prior to the lot expiration date. The following antibodies were used for isPLA: mouse anti-integrin-α8, MAB6194, www.rndsystems.com; rabbit anti-integrin-β1, sc-8978, www.scbt.com; rabbit anti-Mfge8, HPA002807, www.sigmaaldrich.com; rabbit anti-enteric γ-actin, GTX55849, www.genetex.com.

### Confocal microscopy and image acquisition

The slides were examined using an LSM510 Meta (Zeiss, www.zeiss.com) or Fluoview FV1000 confocal microscope (Olympus,www.olympus-lifescience.com) [34]. Confocal micrographs are digital composites of the Z-series of scans (1 µm optical sections of 10 µm thick sections). Settings were fixed at the beginning of both acquisition and analysis steps and were unchanged. Brightness and contrast were slightly adjusted after merging. Final images were constructed using FV10-ASW 2.1 software (Olympus). Each image is representative of labeling experiments from 3 sections from 3 gastric antrum muscles. Scale bars, 10 µm.

### Data and Statistical analysis

Contractile responses were compared by measuring the area under the curve (AUC) of each peak including the contribution of basal tone (integral, grams × seconds) divided by time (seconds), per cross-sectional area (cm^2^) of the smooth muscles, using Acknowledge. The average peak responses (mean (SD)) were calculated using Prism, and significance was determined by *t* test using Prism with *P* < 0.05 considered as significant. Graphs were generated using Prism. The area of the peak chemiluminescence intensity values of the protein bands were calculated by Compass software. The chemiluminescence intensity values of pT696, pT853, and pS19 were divided by the total MYPT1, and LC20 chemiluminescence intensity values from the same sample, respectively, to obtain the ratio of phosphorylated protein to total protein. The ratios were normalized to 1 for unstimulated muscles and all ratios were subsequently analyzed by non-parametric repeated tests of ANOVA using Prism 7.01 software (GraphPad Software,www.graphpad.com), and are expressed as the means ± SD. Student’s t test was used to measure significance and P<0.05 is considered significant. The digital lane views (bitmaps) of the immunodetected protein bands were generated by Compass software, with each lane corresponding to an individual capillary tube. The isPLA figures were created from the digitized data using Adobe Photoshop Version 12.0.3. Graphs were generated using GraphPad/Prism.

### Drugs and reagents

Recombinant human Mfge8 and recombinant human laminin subunit alpha-1 were purchased from R&D Systems, www.rndsystems.com; atropine and tetrodotoxin were obtained from EMD Millipore, www.emdmillipore.com; and MRS2500 was purchased from Tocris Bioscience, www.tocris.com. All other reagents and chemicals purchased were of analytical grade or better.

## Results

### Human gastric antrum muscles express Mfge8, α8 integrin, and β1 integrin

Since Mfge8 and α8 integrin expression in human gastric antrum muscles has not been reported, we examined homogenates of human gastric antrum muscles for Mfge8 and α8 integrin protein expression, along with β1 integrin protein expression. Similar to murine gastric antrum muscles, human gastric antrum muscles express Mfge8 (43kDa), α8 integrin (118kDa), and β1 integrin (89kDa), as shown by the Wes analysis of human gastric antrum muscle lysates in Figure 1.

**Figure 1.**
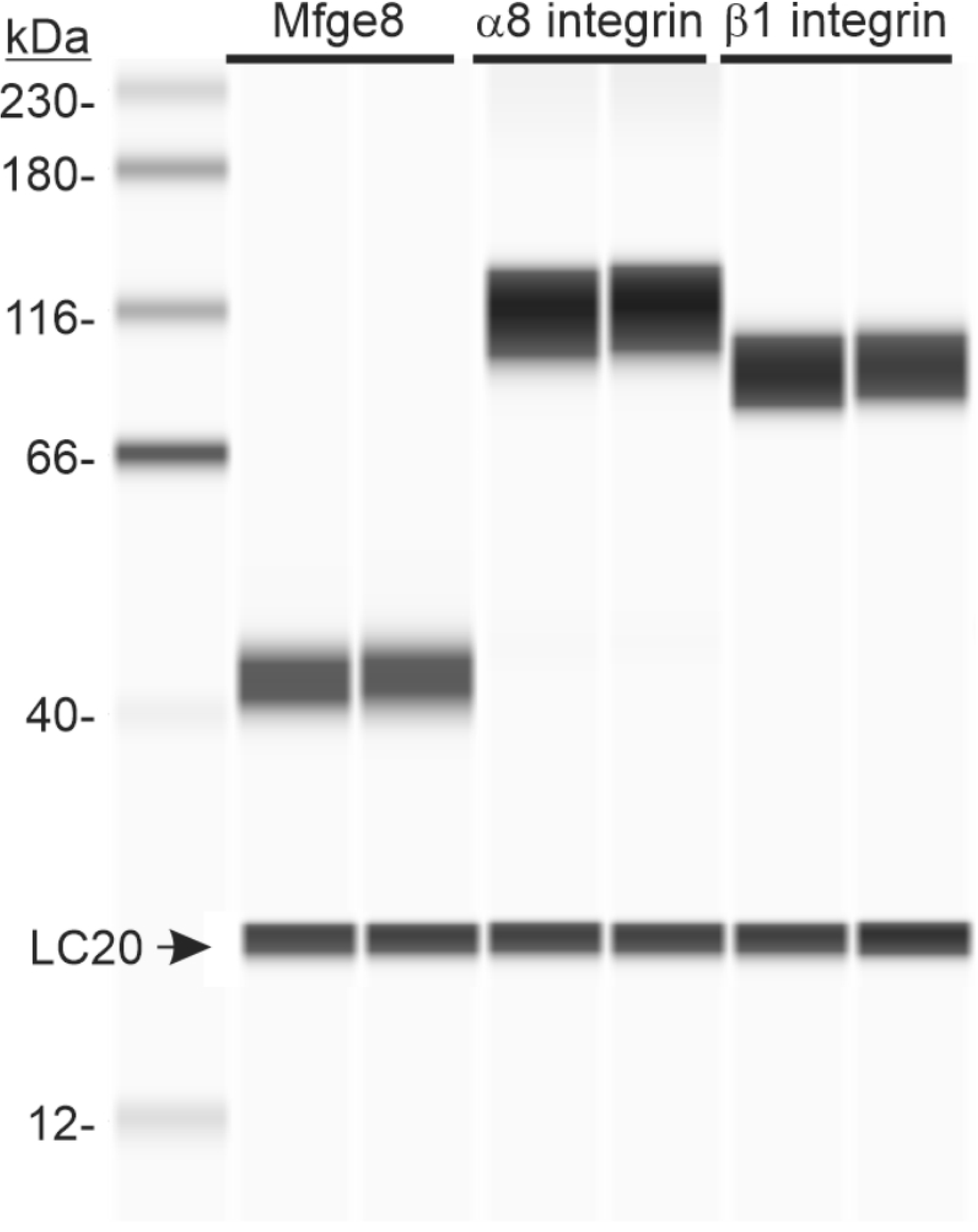
Mfge8, α8 integrin, and β1 integrin are expressed in human gastric antrum smooth muscles. Representative Wes image of Mfge8, α8 integrin, and β1 integrin proteins in gastric antrum smooth muscle by chemiluminescence immunodetection using anti-Mfge8 (100X dilution), α8 integrin (100X dilution), and β1 integrin (100X dilution) antibodies in duplicate as described in the Methods. 5.0µg lysate protein per lane. Anti-LC20 (1:500 dilution) immunodetection was used as the loading control.

### Human gastric antrum muscles contain a8β1 integrin heterodimers

Because it was previously reported by Khalifeh-Soltani *et al*., 2016b that Mfge8 binds to α8 integrin in a8β1 integrin heterodimers in murine gastric antrum muscles, we used in situ PLA to determine whether Mfge8 binds to α8 integrin in a8β1 integrin heterodimers in human gastric antrum muscles. We also immunostained entericactin to localize smooth muscles cells in the antrum smooth muscle sections. The isPLA results and enteric γ-actin immunostaining in Figure 2A show that a8β1 integrin heterodimers are present in human gastric antrum smooth muscles. We then carried out in situ PLA using anti α8 integrin and anti Mfge8 antibodies to determine whether human gastric antrum smooth muscles contain Mfge8 bound to α8 integrin. We also immunostained β1 integrin to localize smooth muscle cell plasma membranes in the antrum smooth muscle sections. The isPLA results and β1 integrin immunostaining in Figure 2B show that Mfge8 is likely bound to α8 integrin in human gastric antrum smooth muscles.

**Figure 2.**
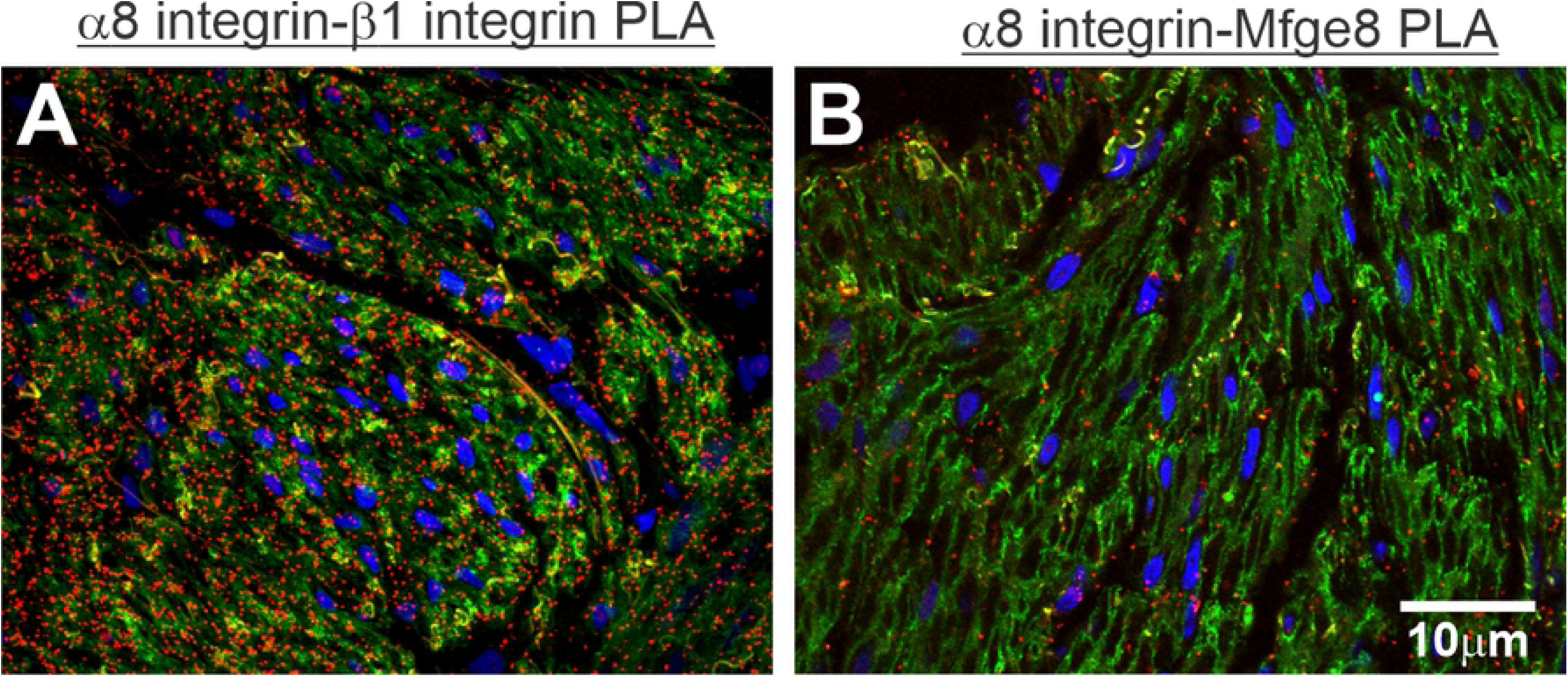
α8β1 integrin heterodimers and Mfge8 interactions with α8 integrin in human gastric antrum smooth muscle shown by in situ PLA. Representative confocal microscopy images from gastric antrum smooth muscle sections. A. Section immunostained with enteric γ-actin (green), and then probed with anti-α8 integrin and β1 integrin antibodies for PLA immunostaining (red spots). B. Section immunostained with β1 integrin (green), and then probed with anti-Mfge8 and α8 integrin antibodies for PLA immunostaining (red spots).

Exogenously added Mfge8 inhibits CCh-evoked contractions of human gastric antrum muscles. We next determined if Mfge8 can regulate human gastric antrum muscle contractile responses. Figure 3 shows the isometric contractile responses of human gastric antrum muscle strips to the cholinergic agonist CCh. CCh at concentrations of 1µM and 5µM dose-dependently increased the force of contractions, as shown in the contractile recordings and the summarized data. After washout of CCh, Mfge8 was added to the myobaths at a concentration of 100µg/ml, and incubated with the muscle strips for 90 minutes. Laminin was added to separate myobaths at a concentration of 100µg/ml, as a negative control integrin RGD-binding protein [36]. As shown in Figs. 3B and 3C, the addition of Mfge8 cause a rapid, but transient contraction of the muscle strips, while laminin had no effect upon addition to the myovbath. As shown in Figs. 3A and 3D, the contractile responses to 5µM CCh 90 minutes after the first 5µM CCh-evoked contraction were unchanged. Similarly, after incubation with laminin for 90 minutes, Figs. 3B and 3E show that the contractile responses of human gastric antrum muscle strips to 5µM CCh were similar to the first 5µM CCh-evoked contraction. In contrast, Figs. 3C and 3F show that compared to the first 5µM CCh-evoked contraction, the contractile response of human gastric antrum muscle strips to 5µM CCh was significantly decreased by incubation with Mfge8 for 90 minutes. In addition, Figs. 3C and 3F show that the contractile responses of the muscle strips to 5µM CCh recovered following washout of Mfge8, as indicated by the increase in the AUC.

**Figure 3.**
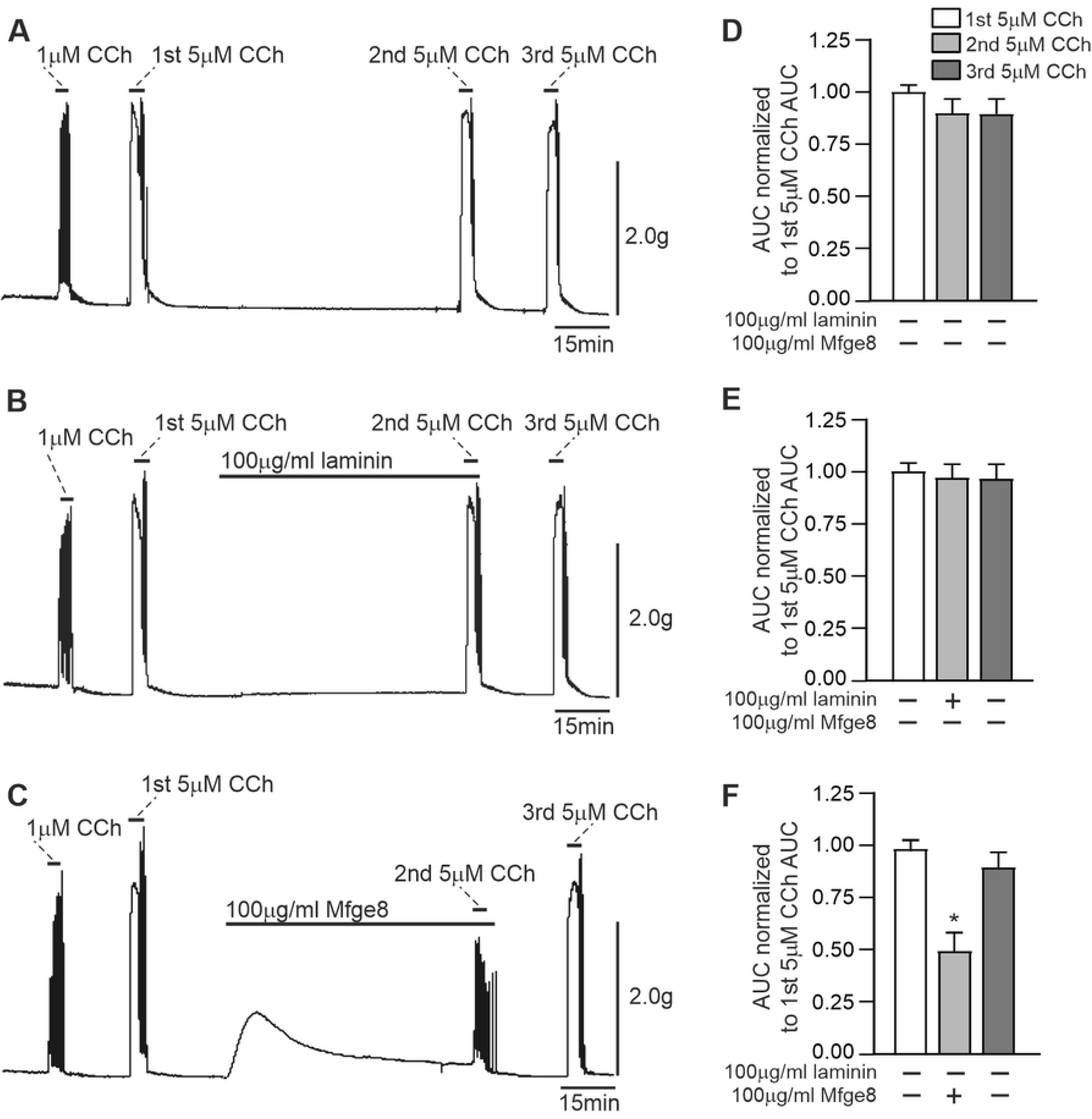
Exogenously added Mfge8 inhibits CCh-evoked contractions of human gastric antrum smooth muscle. Representative tension recordings of the contractile responses to 5µM CCh alone (A), or in the presence of 100µg/ml laminin (B), or 100µg/ml Mfge8 (C). Summarized data of the areas under the curve of each contractile response (D,E,F). (n= 6; 2 muscle strips from 3 gastric antrums; *P<0.05).

### Exogenously added Mfge8 inhibits MYPT1 and LC20 phosphorylation in human gastric antrum muscles

It was previously determined that Mfge8 inhibits murine gastric antrum muscle contractions by inhibiting MYPT1 pT696 phosphorylation, resulting in decreased LC20 phosphorylation [27]. Since we found that Mfge8 inhibits human gastric antrum muscle contractions, we examined whether CCh-evoked MYPT1 and LC20 phosphorylation are inhibited by Mfge8. As shown in Figs. 4A and 4B, 5 min treatment with 5µM CCh increased MYPT1 T696 and T853 phosphorylation. Incubation with laminin for 90 minutes had no effect on the CCh-evoked increase in MYPT1 T853 phosphorylation and did not affect T696 phosphorylation. However, Figs. 4A and 4B show that the CCh-evoked increase in MYPT1 T853 phosphorylation was significantly inhibited by incubation with Mfge8 for 90 minutes, and MYPT1 pT696 phosphorylation was reduced. Figures 4C and 4D show that LC20 S19 phosphorylation was consistently increased by CCh treatment, but this increase was not statistically significant. Laminin had no effect on the increase in LC20 S19 phosphorylation (Figs. 4C, 4D). In contrast, the CCh-evoked increase in LC20 S19 phosphorylation was inhibited by incubation with Mfge8 for 90 minutes, but this decrease was not statistically significant.

**Figure 4.**
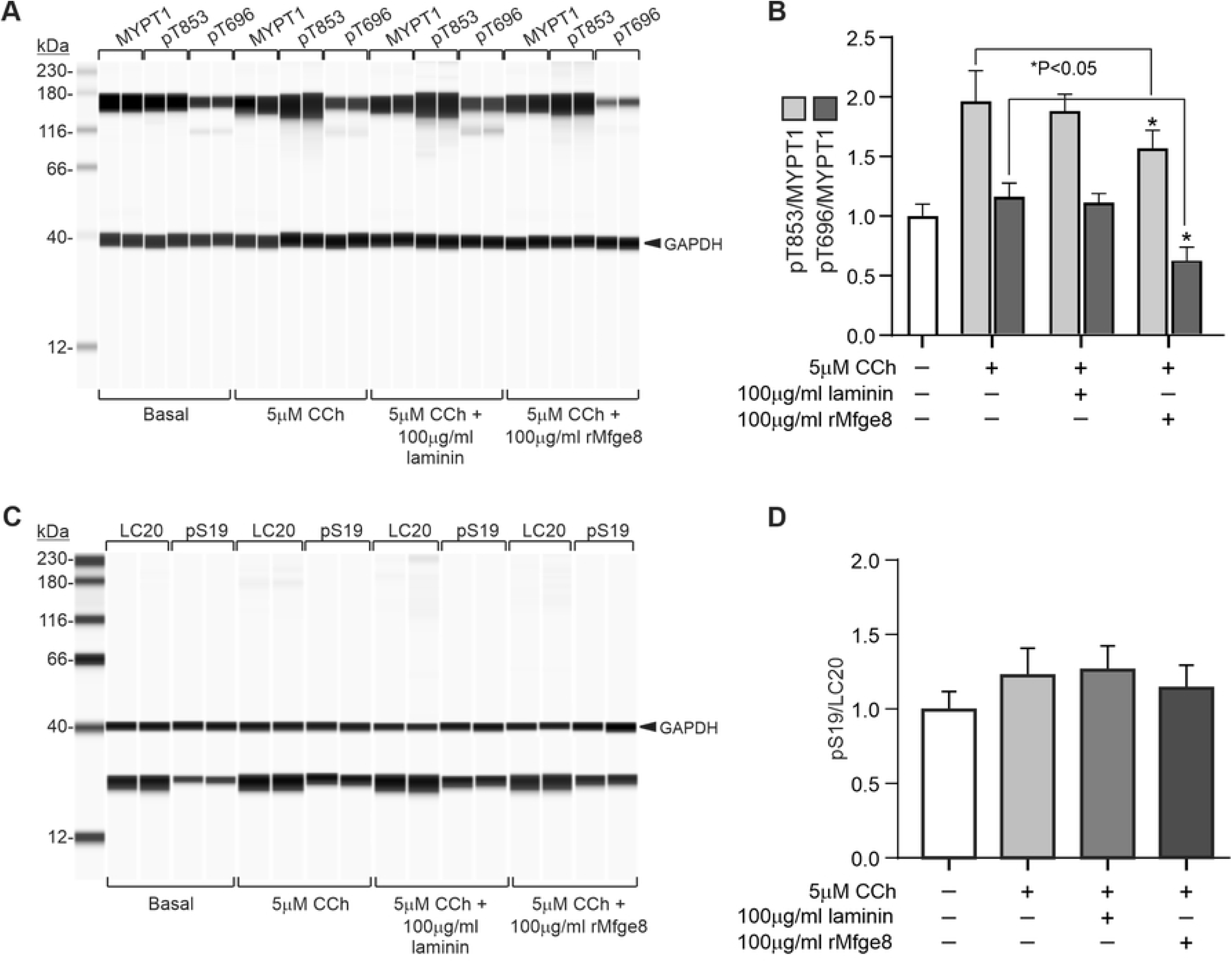
Exogenously added Mfge8 inhibits CCh-evoked phosphorylation of MYPT1 and LC20 in human gastric antrum smooth muscles. A. Representative Wes analysis of MYPT1 T853 and T696 phosphorylation by 5µM CCh alone, or in the presence of 100µg/ml laminin, or 100µg/ml Mfge8. B. Summary of the effects of 5µM CCh alone, or in the presence of 100µg/ml laminin, or 100µg/ml Mfge8 on MYPT1 T853 and T696 phosphorylation. C. Representative Wes analysis of LC20 S19 phosphorylation by 5µM CCh alone, or in the presence of 100µg/ml laminin, or 100µg/ml Mfge8. D. Summary of the effects of 5µM CCh alone, or in the presence of 100µg/ml laminin, or 100µg/ml Mfge8 on LC20 S19 phosphorylation. GAPDH immunodetection was used as the loading control. (n= 6; 2 muscle strips from 3 gastric antrums).

### Exogenously added Mfge8 inhibits contractions of human gastric antrum muscles evoked by electrical field stimulation (EFS)

Having found that Mfge8 reduces MYPT1 and LC20 phosphorylation and inhibits CCh-evoked contractions, we then determined if Mfge8 inhibits the contractile responses to endogenous cholinergic motor neurotransmission. The isometric contractile responses of human gastric antrum smooth muscles to 5Hz, 10Hz, and 20Hz EFS were obtained in the presence of LNNA and MRS2500 to block nitrergic and purinergic neurotransmission. Figure 5A shows that contractile responses were increased in a frequency dependent manner, and were completely blocked by atropine. As shown in Fig. 5B, the contractile responses to 5Hz, 10Hz, and 20Hz EFS 90 minutes after the first set of EFS-evoked contractions were unchanged. Mfge8 was added to the myobaths at a concentration of 100µg/ml, and incubated with the muscle strips for 90 minutes. Figure 5C shows that compared to the first set of EFS-evoked contractions, the EFS-evoked contractile responses to 5Hz, 10Hz, and 20Hz EFS were significantly inhibited by incubation with 100µg/ml Mfge8 for 90 minutes.

**Figure 5.**
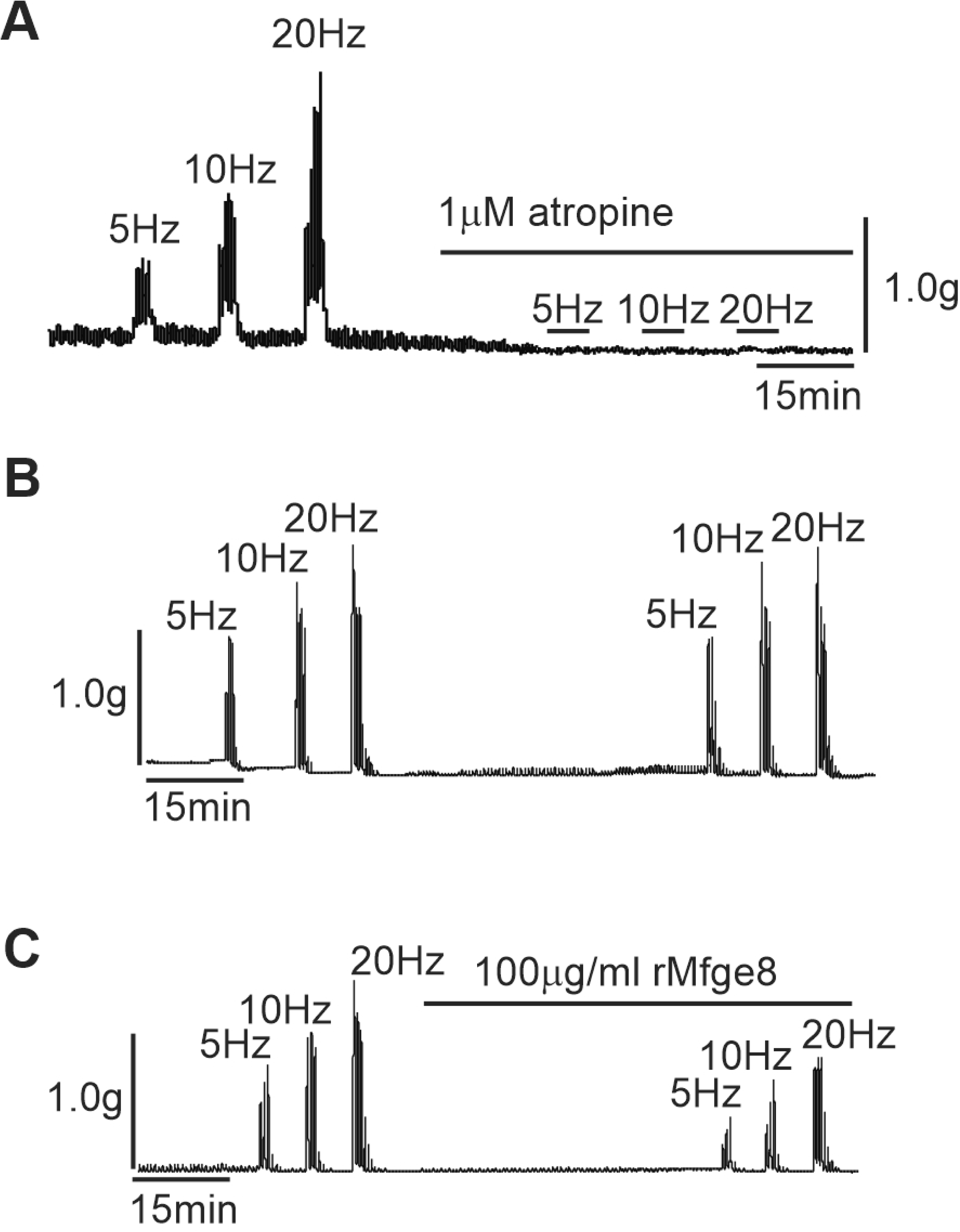
Exogenously added Mfge8 inhibits EFS-evoked cholinergic contractions of human gastric antrum smooth muscle. A. Representative tension recording of the contractile responses to 5Hz, 10Hz, 20Hz alone, or in the presence of 1µM atropine. B. Representative tension recording of the contractile responses to 5Hz, 10Hz, 20Hz alone. C. Representative tension recording of the contractile responses to 5Hz, 10Hz, 20Hz alone, or in the presence of 100µg/ml Mfge8. (n= 3; 1 muscle strip from 3 gastric antrums).

## Discussion

It was previously reported by Khalifeh-Solani et al. that in mice, Mfge8 inhibits antral muscle contractions and slows gastrointestinal motility by specifically binding to αβ integrin in α8β1 integrin heterodimers, resulting in reduced phosphorylation of the inhibitory MYPT1 subunit of MLCP, and consequentially reduced LC20 phosphorylation [27]. In addition, either smooth muscle-specific deletion of Mfge8 or α8 resulted in an increase in gastric antral contractile force, more rapid gastric emptying, and faster small intestinal transit times [27]. These findings revealed a novel inhibitory mechanism regulating gastric antrum function, raising the question as to whether a similar mechanism is involved in regulating human gastric antrum smooth muscle contractile responses. The expression of Mfge8 or α8 integrin in human gastric antrum muscles has not been described previously, thus in this study we determined that both Mfge8 and α8β1 integrin heterodimers are present in human gastric antrum muscles, and that Mfge8 is bound to α8β1 integrin heterodimers. We also show that exogenously added Mfgfe8 inhibits the contractile responses of human gastric antrum muscles to exogenous and endogenous cholinergic stimuli. This inhibition of contraction was accompanied by inhibition of MYPT1 and LC20 phosphorylation, supporting a novel role for α8β1 integrins and Mfge8 in regulating human gastric motility by attenuating MYPT1 phosphorylation. The findings that both Mfge8 and α8β1 integrin heterodimers are present in human gastric antrum muscles, suggest that Mfge8 is involved in the regulation of human gastric antrum muscle mechanical responses. We used in situ PLA to demonstrate the interaction between Mfge8 and α8 integrin. We were not able to examine the effects of abrogating the binding of Mfge8 to α8β1 integrins because there is no inhibitor of Mfge8 binding to α8β1 integrins available. However, adding Mfge8 protein to the muscle strips in the myobaths significantly inhibited the contractile responses to the cholinergic agonist CCh or to EFS-evoked cholinergic neurotransmission. These findings suggest that there are α8β1 integrins not occupied by Mfge8, and that increases in Mfge8 could further inhibit gastric antrum muscle contraction.

Mfge8 (originally named lactadherin) was first identified in breast milk, having antimicrobial and antiviral effects, and playing an important role in immune defense as a secreted immune system molecule [37, 38]. Mfge8 is now known to be a ubiquitously expressed multifunctional protein belonging to the family of secreted integrin-binding glycoproteins containing the RGD integrin-binding motif [39]. The most well known role for α8β1 is in kidney morphogenesis where deletion of α8 integrin leads to impaired recruitment of mesenchymal cells into epithelial structures[40, 41]. α8 integrin is a member of the RGD-binding integrin family that is prominently expressed in smooth muscle coupled to β1 integrin [42-44]. Previous work has shown the expression of α8 integrin in both vascular and visceral smooth muscle, as well as the muscularis mucosa of the GI tract [42]. In vitro studies suggest that α8 promotes smooth muscle differentiation, and maintains vascular smooth muscle in a differentiated, contractile, non-migratory phenotype [43, 45]. Mfge8 and α8 integrin also modulate smooth muscle contractile force. In Mfge8^-/-^ mice, or α8 integrin^-/-^ mice, airway and jejunal smooth muscle contraction are enhanced in response to contractile agonists after these muscle beds have been exposed to inflammatory cytokines but not under basal conditions [27, 46, 47]. Whether the origin of Mfge8 in gastric muscles is from circulating Mfge8 or locally secreted is unclear. Mfge8 can reach the gastric antrum smooth muscle layer by oral gavage, but it is not clear how Mfge8 reaches the gastric antrum smooth muscle layer, or how widespread the distribution of Mfge8 is after oral administration [27]. Determining the source of Mfge8 present in gastric muscle tissues is an important issue to address in future studies of gastric motility regulatory mechanisms.

In summary, in this study we report that the secreted protein Mfge8 inhibits the contractile responses of human gastric antrum muscles to cholinergic stimuli by reducing the inhibitory phosphorylation of the MYPT1 subunit of MLCP, resulting in reduced LC20 phosphorylation. We found that endogenous Mfge8 is bound to its receptor, α8β1 integrin, in human gastric antrum muscles, suggesting that human gastric antrum muscle mechanical responses are regulated by Mfge8. These findings, and the findings of Khalifeh-Soltani et al. 2016, reveal an additional pathway regulating the contractile responses of smooth muscles. Elevations in cytosolic Ca2+ directly promote smooth muscle contraction by Ca^2+^/calmodulin activation of MLCK and phosphorylation of LC20 [9]. Rho kinase and PKC activities contribute to MLCK activity by phosphorylating the regulatory subunits of MLCP to promote LC20 phosphorylation and increase the myofilament sensitivity to Ca2+ [48]. In addition, a number of studies have provided evidence that dynamic changes to the actin cytoskeleton play an important role in smooth muscle contraction [49, 50]. This remodeling process is thought to facilitate the polymerization of cortical cytoskeletal actin filaments and increase the stability of focal adhesions in the membrane, allowing for the force generated by myofilament activation to be transmitted to the connective tissue of the extracellular matrix [51, 52]. Tyrosine phosphorylation of protein tyrosine kinase 2 β (Pyk2) and focal adhesion kinase (FAK), along with the recruitment of other integrin-associated proteins to focal adhesions, occurs during contraction and force development [53]. In addition, we found that FAK also promotes gastric smooth muscle contraction by activation of the PKC-CPI-17 Ca^2+^ sensitization pathway [33]. The regulation of gastric antrum smooth muscles by Mfge8 and α8 integrin opposes the prokinetic actions of MLCK activation, MLCP inhibition, and cytoskeletal remodeling. In this regard, Mfge8 α8 integrin signaling seems to function as a brake on gastric antrum mechanical activities. Further studies of the role of Mfge8 and α8 integrin in regulating gastric antrum function will likely reveal additional novel aspects of gastric smooth muscle motility mechanisms.

## Acknowledgements

The research reported in this publication was supported by a National Institute of Diabetes and Digestive and Kidney Diseases Diabetic Complications Consortium (DiaComp, http://www.diacomp.org) Grant DK076169, and a Takeda Pharmaceuticals Innovation Center Grant to BAP, and by a Mick Hitchcock Graduate Student Scholarship to YX. We thank Drew Syder and Jill Wykosky of the Takeda GI Drug Discovery Unit for their enthusiastic guidance and expert advice during the course of this study.

## References

1. Janssen P, Vanden Berghe P, Verschueren S, Lehmann A, Depoortere I, Tack J. Review article: the role of gastric motility in the control of food intake. Aliment Pharmacol Ther 2011;33(8):880-894. pmid: 21342212.

2. Kong F, Singh RP. Disintegration of solid foods in human stomach. J Food Sci. 2008;73(5):R67-R80. pmid: 18577009.

3. Tack J, Janssen P. Gastroduodenal motility. Curr Opin Gastroenterol. 2010;26:647-655. pmid: 20838344.

4. Goyal RK, Guo Y, Mashimo H. Advances in the physiology of gastric emptying. Neurogastroenterol Motil. 2019;31(4):e13546. pmid: 30740834.

5. Tack J, Masuy I, Van Den Houte K, Wauters L, Schol J, Vanuytsel T, et al. Drugs under development for the treatment of functional dyspepsia and related disorders. Expert Opin Investig Drugs. 2019;28(10):871-889. pmid: 31566013.

6. Sanders KM, Koh SD, Ro S, Ward SM. Regulation of gastrointestinal motility--insights from smooth muscle biology. Nat Rev Gastroenterol Hepatol. 2012;9(11):633-645. pmid: 22965426.

7. Zhang RX, Wang XY, Chen D, Huizinga JD. Role of interstitial cells of Cajal in the generation and modulation of motor activity induced by cholinergic neurotransmission in the stomach. Neurogastroenterol Motil. 2011;23(9):e356-e371. pmid: 21781228.

8. He WQ, Peng YJ, Zhang WC, Lv N, Tang J, Chen C, et al. Myosin light chain kinase is central to smooth muscle contraction and required for gastrointestinal motility in mice. Gastroenterol. 2008;135(2):610-620. pmid: 18586037.

9. Somlyo AP, Somlyo AV. Ca2+ sensitivity of smooth muscle and nonmuscle myosin II: Modulated by G proteins, kinases, and myosin phosphatase. Physiol Rev. 2003;83(4):1325-1358. pmid: 14506307.

10. Somlyo AP, Himpens B. Cell calcium and its regulation in smooth muscle. FASEB J. 1989;3:2266-2276. pmid: 2506092.

11. Somlyo AP, Somlyo AV. Electron probe analysis of calcium content and movements in sarcoplasmic reticulum, endoplasmic reticulum, mitochondria, and cytoplasm. J Cardiovasc Pharmacol. 1986;8 Suppl 8:S42-S47. pmid: 2433524.

12. Alessi D, MacDougall LK, Sola MM, Ikebe M, Cohen P. The control of protein phosphatase-1 by targeting subunits. The major myosin phosphatase in avian smooth muscle is a novel form of protein phosphatase-1. Eur J Biochem. 1992;210:1023-1035. pmid: 1336455

13. Paul RJ, Shull GE, Kranias EG. The sarcoplasmic reticulum and smooth muscle function: evidence from transgenic mice. Novartis Found Symp. 2002;246:228-238. pmid: 12164311

14. Feng J, Ito M, Ichikawa K, Isaka N, Nishikawa M, Hartshorne DJ, et al. Inhibitory phosphorylation site for Rho-associated kinase on smooth muscle myosin phosphatase. J Biol Chem. 1999;274(52):37385-37390. pmid: 10601309.

15. Ito M, Nakano T, Erdodi F, Hartshorne D. Myosin phosphatase: Structure, regulation and function. Mol Cell Biochem. 2004;259:197-209. pmid:15124925.

16. Kitazawa T, Eto M, Woodsome TP, Khalequzzaman M. Phosphorylation of the myosin phosphatase targeting subunit and CPI-17 during Ca2+ sensitization in rabbit smooth muscle. J Physiol. 2003;546(3):879-889. pmid: 2342583.

17. Eto M, Ohmori T, Suzuki M, Furuya K, Morita F. A novel protein phosphatase-1 inhibitory protein potentiated by protein kinase C. Isolation from porcine aorta media and characterization. J Biochem. 1995;118(6):1104-1107. pmid: 8720121.

18. Hayashi Y, Senba S, Yazawa M, Brautigan DL, Eto M. Defining the structural determinants and a potential mechanism for inhibition of myosin phosphatase by the protein kinase C-potentiated inhibitor protein of 17 kDa. J Biol Chem. 2001;276(43):39858-39863. pmid: 11517233.

19. Grassie ME, Moffat LD, Walsh MP, MacDonald JA. The myosin phosphatase targeting protein (MYPT) family: A regulated mechanism for achieving substrate specificity of the catalytic subunit of protein phosphatase type 1δ. Arch Biochem Biophys. 2011;510(2):147-159. pmid: 21291858.

20. Matsumura F, Hartshorne DJ. Myosin phosphatase target subunit: Many roles in cell function. Biochem Biophys Res Comm. 2008;369(1):149-156. pmid: 18155661

21. Velasco G, Armstrong C, Morrice N, Frame S, Cohen P. Phosphorylation of the regulatory subunit of smooth muscle protein phosphatase 1δ at Thr850 induces its dissociation from myosin. FEBS Lett. 2002;527:101-104. pmid: 12220642.

22. Chen CP, Chen X, Qiao YN, Wang P, He WQ, Zhang CH, et al. In vivo roles for myosin phosphatase targeting subunit-1 phosphorylation sites T694 and T852 in bladder smooth muscle contraction. J Physiol. 2015;593(3):681-700. pmid: 25433069

23. He W-Q, Qiao Y-N, Peng Y-J, Zha J-M, Zhang C-H, Chen C, et al. Altered contractile phenotypes of intestinal smooth muscle in mice deficient in myosin phosphatase target subunit 1. Gastroenterol. 2013;144(7):1456-1465. pmid: 23499953

24. Gao N, Huang J, He W, Zhu M, Kamm KE, Stull JT. Signaling through myosin light chain kinase in smooth muscles. J Biol Chem. 2013. 288(11):7596-605. pmid: 23362260

25. Kitazawa T, Gaylinn BD, Denney GH, Somlyo AP. G-protein-mediated Ca2+ sensitization of smooth muscle contraction through myosin light chain phosphorylation. J Biol Chem. 1991;266(3):1708-1715. pmid: 1671041

26. Mizuno Y, Isotani E, Huang J, Ding H, Stull JT, Kamm KE. Myosin light chain kinase activation and calcium sensitization in smooth muscle in vivo. Am J Physiol - Cell Physiology. 2008;295(2):C358-C364. pmid: 18524939

27. Khalifeh-Soltani A, Ha A, Podolsky MJ, McCarthy DA, McKleroy W, Azary S, et al. α8β1 integrin regulates nutrient absorption through an Mfge8-PTEN dependent mechanism. Elife. 2016;5. pmid: 27092791.

28. Bhetwal BP, An CL, Fisher SA, Perrino BA. Regulation of basal LC20 phosphorylation by MYPT1 and CPI-17 in murine gastric antrum, gastric fundus, and proximal colon smooth muscles. Neurogastroenterol Motil. 2011;23(10):e425-e436. pmid: 21883701

29. Bhetwal BP, An CL, Baker SA, Lyon KL, Perrino BA. Impaired contractile responses and altered expression and phosphorylation of Ca2+ sensitization proteins in gastric antrum smooth muscles from ob/ob mice. J Muscle Res Cell Motil. 2013;34(2):137-149. pmid: 23576331

30. Bhetwal BP, Sanders KM, An C, Trappanese DM, Moreland RS, Perrino BA. Ca2+ sensitization pathways accessed by cholinergic neurotransmission in the murine gastric fundus. J Physiol (Editors’ Choice). 2013;591(Pt 12):2971-2986. pmid: 23613531

31. Rhee PL, Lee JY, Son HJ, Kim JJ, Rhee JC, Kim S, et al. Analysis of pacemaker activity in the human stomach. J Physiol. 2011;589(Pt 24):6105-6118. pmid: 22005683.

32. Li W, Sasse KC, Bayguinov Y, Ward SM, Perrino BA. Contractile protein expression and phosphorylation and contractility of gastric smooth muscles from obese patients and patients with obesity and diabetes. J Diabetes Res. 2018:8743874. pmid: 29955616.

33. Xie Y, Han KH, Grainger N, Li W, Corrigan RD, Perrino BA. A role for focal adhesion kinase in facilitating the contractile responses of murine gastric fundus smooth muscles. J Physiol. 2018;596(11):2131-2146. pmid: 29528115.

34. Xie Y, Perrino BA. Quantitative in situ proximity ligation assays examining protein interactions and phosphorylation during smooth muscle contractions. Anal Biochem. 2019;577:1-13. pmid: 30981700.

35. Ulke-Lemée A, Turner SR, MacDonald JA. In situ analysis of smoothelin-like 1 and calmodulin interactions in smooth muscle cells by proximity ligation. J Cell Biochem. 2015;116(11):2667-2675. pmid: 25923522.

36. Zheng Y, Leftheris K. Insights into protein-ligand interactions in integrin complexes: Advances in structure determinations. J Med Chem. 2020;63(11):5675-5696. pmid: 31999923.

37. Atabai K, Fernandez R, Huang X, Ueki I, Kline A, Li Y, et al. Mfge8 is critical for mammary gland remodeling during involution. Mol Biol Cell. 2005;16(12):5528-5537. pmid: 16195353.

38. Stubbs JD, Lekutis C, Singer KL, Bui A, Yuzuki D, Srinivasan U, et al. cDNA cloning of a mouse mammary epithelial cell surface protein reveals the existence of epidermal growth factor-like domains linked to factor VIII-like sequences. Proc Natl Acad Sci USA. 1990;87(21):8417-8421. pmid: 2122462.

39. Raymond A, Ensslin MA, Shur BD. SED1/MFG-E8: a bi-motif protein that orchestrates diverse cellular interactions. J Cell Biochem. 2009;106(6):957-966. pmid: 19204935.

40. Humbert C, Silbermann F, Morar B, Parisot M, Zarhrate M, Masson C, et al. Integrin alpha 8 recessive mutations are responsible for bilateral renal agenesis in humans. Am J Hum Genet. 2014;94(2):288-294. pmid: 24439109.

41. Müller U, Wang D, Denda S, Meneses JJ, Pedersen RA, Reichardt LF. Integrin alpha8beta1 is critically important for epithelial-mesenchymal interactions during kidney morphogenesis. Cell. 1997;88(5):603-613. pmid: 9054500.

42. Schnapp LM, Hatch N, Ramos DM, Klimanskaya IV, Sheppard D, Pytela R. The human integrin alpha 8 beta 1 functions as a receptor for tenascin, fibronectin, and vitronectin. J Biol Chem. 1995;270(39):23196-23202. pmid: 7559467.

43. Zargham R, Thibault G. Alpha 8 integrin expression is required for maintenance of the smooth muscle cell differentiated phenotype. Cardiovasc Res. 2006;71(1):170-178. pmid: 16603140.

44. Zargham R, Touyz RM, Thibault G. alpha 8 Integrin overexpression in de-differentiated vascular smooth muscle cells attenuates migratory activity and restores the characteristics of the differentiated phenotype. Atherosclerosis. 2007;195(2):303-312. pmid: 17275006.

45. Zhang MJ, Zhou Y, Chen L, Wang YQ, Wang X, Pi Y, et al. An overview of potential molecular mechanisms involved in VSMC phenotypic modulation. Histochem Cell Biol. 2016;145(2):119-130. pmid: 26708152.

46. Khalifeh-Soltani A, Gupta D, Ha A, Podolsky MJ, Datta R, Atabai K. The Mfge8-α8β1-PTEN pathway regulates airway smooth muscle contraction in allergic inflammation. Faseb J. 2018:fj201800109R. pmid: 29763381.

47. Kudo M, Khalifeh Soltani SM, Sakuma SA, McKleroy W, Lee TH, Woodruff PG, et al. Mfge8 suppresses airway hyperresponsiveness in asthma by regulating smooth muscle contraction. Proc Natl Acad Sci USA. 2013;110(2):660-665. pmid: 23269839.

48. Perrino BA. Calcium Sensitization Mechanisms in Gastrointestinal Smooth Muscles. J Neurogastroenterol Motil. 2016. 22(2):213-25. pmid: 26701920

49. Mehta D, Gunst SJ. Actin polymerization stimulated by contractile activation regulates f force development in canine tracheal smooth muscle. J Physiol. 1999;519 Pt 3(Pt 3):829-840. pmid: 10457094.

50. Zhang W, Bhetwal BP, Gunst SJ. Rho kinase collaborates with p21-activated kinase to regulate actin polymerization and contraction in airway smooth muscle. J Physiol. 2018;596(16):3617-35. pmid: 29746010.

51. Mills RD, Mita M, Nakagawa J, Shoji M, Sutherland C, Walsh MP. A role for the tyrosine kinase Pyk2 in depolarization-induced contraction of vascular smooth muscle. J Biol Chem. 2015;290(14):8677-8692. pmid: 25713079.

52. Zheng C, Xing Z, Bian ZC, Guo C, Akbay A, Warner L, et al. Differential regulation of Pyk2 and focal adhesion kinase (FAK). The C-terminal domain of FAK confers response to cell adhesion. J Biol Chem. 1998;273(4):2384-389. pmid: 9442086.

53. Gerthoffer WT, Gunst SJ. Invited Review: Focal adhesion and small heat shock proteins in the regulation of actin remodeling and contractility in smooth muscle. J Appl Physiol. 2001;91(2):963-972. pmid: 11457815.

